# Cortex-wide topography of 1/f-exponent in Parkinson’s disease

**DOI:** 10.1101/2023.01.19.524792

**Authors:** Pascal Helson, Daniel Lundqvist, Per Svenningsson, Mikkel C. Vinding, Arvind Kumar

## Abstract

Parkinson’s Disease causes progressive and debilitating changes to the brain as well as to the mind. While the diagnostic hallmark features are the characteristic movement-related symptoms, the disease also causes decline in sensory processing, cognitive, emotional performance and most patients develop dementia over time. The extent of symptoms and the brain-wide projections of neuromodulators such as dopamine suggest that many brain regions are simultaneously affected in Parkinson’s disease. To characterise such disease-related and brain-wide changes in neuronal function, we performed a source level analysis of resting state magnetoencephalogram (MEG) from two groups: Parkinson’s disease patients and healthy controls. Besides standard spectral analysis, we quantified the aperiodic component of the neural activity by fitting a power law (*κ/f* ^λ^) to the MEG spectrum and then studied its relationship with age and UPDRS. Consistent with previous results, the most significant spectral changes were observed in the high theta/low alpha band (7-10 Hz) in all brain regions. Furthermore, analysis of the aperiodic part of the spectrum showed that, in all but frontal regions, λ was significantly larger in Parkinson’s disease patients than in control subjects. Our results indicate for the first time that Parkinson’s disease is associated with significant changes in population activity across the whole neocortex. Surprisingly, even early sensory areas showed a significantly larger λ in patients than in healthy controls. Moreover, λ was not affected by the L-dopa medication. Finally, λ was positively correlated with patient age but not with UPDRS-III (summary measure of motor symptoms’ clinical rating). Because λ is closely associated excitation-inhibition balance, our results propose new hypotheses about manifestation of Parkinson’s disease in cortical networks.

## Introduction

In Parkinson’s Disease (PD), the progressive loss of the dopaminergic cells not only depletes the neuromodulator dopamine but also alters the dynamics of other key neuromodulators such as serotonin, noradrenaline and acetylcholine (see the review by McGregor and Nelson^1^). Given the widespread prevalence of neuromodulators in the neocortex, it is expected that the neocortical neural circuits dynamics and function would also be affected in PD. That is, the signature of PD related dysfunction should also be visible in the neuronal activity recorded from neocortical regions.

Consistent with this, analyses of EEG and MEG have revealed several changes in the population activity of different neocortical regions (see reviews by Geraedts et al.^2^ and Boon et al.^3^). While the results are diverse given the heterogeneity of the patients, it is commonly observed that the low frequency – delta to low-alpha band – power increases whereas high-alpha to gamma band power decreases.^4^

This kind of spectral slowing was shown to be correlated with motor and cognitive symptoms.^5^ Such spectral slowing has been observed in the earliest stages of the disease (e.g. in the posterior cortex regions^6^), hence demonstrating that it is not an effect of dopamine medication. Moreover, dopamine replacement therapy hardly reverses the spectral slowing, especially in the more advanced PD patients.^6^

Spectral power alterations are also associated with changes in functional connectivity (FC) in PD patients. Throughout the disease, the low-alpha band FC decreases after an initial increase.^7,8^ Increase in beta band synchrony is also commonly observed in the basal ganglia as well as in the cortico-basal ganglia loops (see the review by Hammond et al.^9^).

Thus, previous work on analysis of EEG/MEG has been largely focused on oscillatory activity in different frequency bands. Besides oscillations, the aperiodic part of the population activity can also be informative about the underlying network dysfunction and relative excitation-inhibition balance.^10^ To the best of our knowledge, the aperiodic part of the EEG/MEG activity in PD was characterised in three studies. First, Vinding et al.^11^ reported steeper power law decay of MEG power in sensorimotor regions of PD patients. In a more recent study, Wiesman et al.^12^ estimated the aperiodic activity on four different frequency bands showing a neurophysiological slowing – decrease through frequency of the λ deviation from HC. Finally, Wang et al.^13^ studied low-spatial resolution EEG from PD-ON and OFF medication showing that there is a significant increase of the offset and exponent power law parameters from OFF to ON medication in some of the sensors. However, thus far it has remained unclear how the spatial distribution of aperiodic component of the population activity is altered by chronic dopamine depletion and dopamine replacement therapy in human patients.

Therefore, we studied the spatial distribution of spectral peaks and aperiodic component of the activity from neural populations measured with MEG in PD patients in their on and off medication states. To this end, MEG acquired using 306 sensors were pre-processed to obtain 44 sources’ activity distributed all over the neocortex according to the HCP-MMP1 atlas. We found that indeed there is a slowing in the MEG spectrum even when the spectral peaks were identified without classical frequency band definition. Analysis of the power-law exponent (λ) of MEG spectrum revealed a significant increase in the λ in sensory and motor regions in PD patients compared to the healthy controls. In fact, λ showed a spatial positive gradient from anterior to posterior brain regions in PD patients. Surprisingly, the frontal regions which receive most of the dopaminergic projections did not show a significant difference in λ. L-dopa administration did not affect the spatial distribution of λ. However, λ changes were correlated to the patients’ age but not to their UPDRS-III score (the clinical rating of motor symptoms). That is, neocortical activity dynamics is more vulnerable to chronic dopamine changes in older PD patients than in younger patients. Moreover, our results reveal aspects of neocortical population activity which are not affected by the L-dopa therapy. Because 1/f-exponent (λ) can be linked to excitation inhibition (EI) balance, our analysis suggests new testable hypotheses about the PD related changes in the neocortex.

## Materials and methods

### MEG data and its pre-processing

Here we analysed the resting state (eyes opened) MEG recorded from 17 PD patients (age 41-85; five female) and 20 age matched healthy controls (HC; age 54-76; eight female). The data was acquired at the Swedish National Facility for MEG (NATMEG, https://natmeg.se/) using the Elekta Neuromag TRIUX 306-channel MEG system. The study was approved by the regional ethics committee (Etikpöingsnöamden Stockholm, DNR: 2016/911-31/1) and followed the Declaration of Helsinki. All participants gave written informed consent before participating.

For more details of the experimental protocol please refer to Vinding et al.^14^ study. Briefly, the MEG was acquired at a sampling rate of 1 kHz with an online 0.1 Hz high-pass filter and 330 Hz low-pass filter. Each subject was recorded twice. MEG from PD patients was recorded in OFF and ON medication states. In the OFF medication condition, PD patients were off their dopamine replacement medications (Levodopa) for at least 12 hours. After the first recording session, patients took their medication and MEG was recorded for the second time one hour after the medication intake. HC subjects were also recorded twice at an interval of 1 h.

Each recording epoch was of 8 minutes duration. Data were down-sampled to a sampling frequency of 200 Hz and a low pass filtered at 45 Hz was applied. We pre-processed the data to remove non-neural noise and eye movement artifacts using ICA and data from electro-oculogram or electrocardiogram. We then used the dynamic statistical parametric mapping (dSPM^15^) for source reconstruction (implemented in MNE-Python^16^), followed by a labelling using HCP MM1 atlas^17^ which resulted in 44 brain regions (BR) signals as shown in Figure 1.

**Figure 1:**
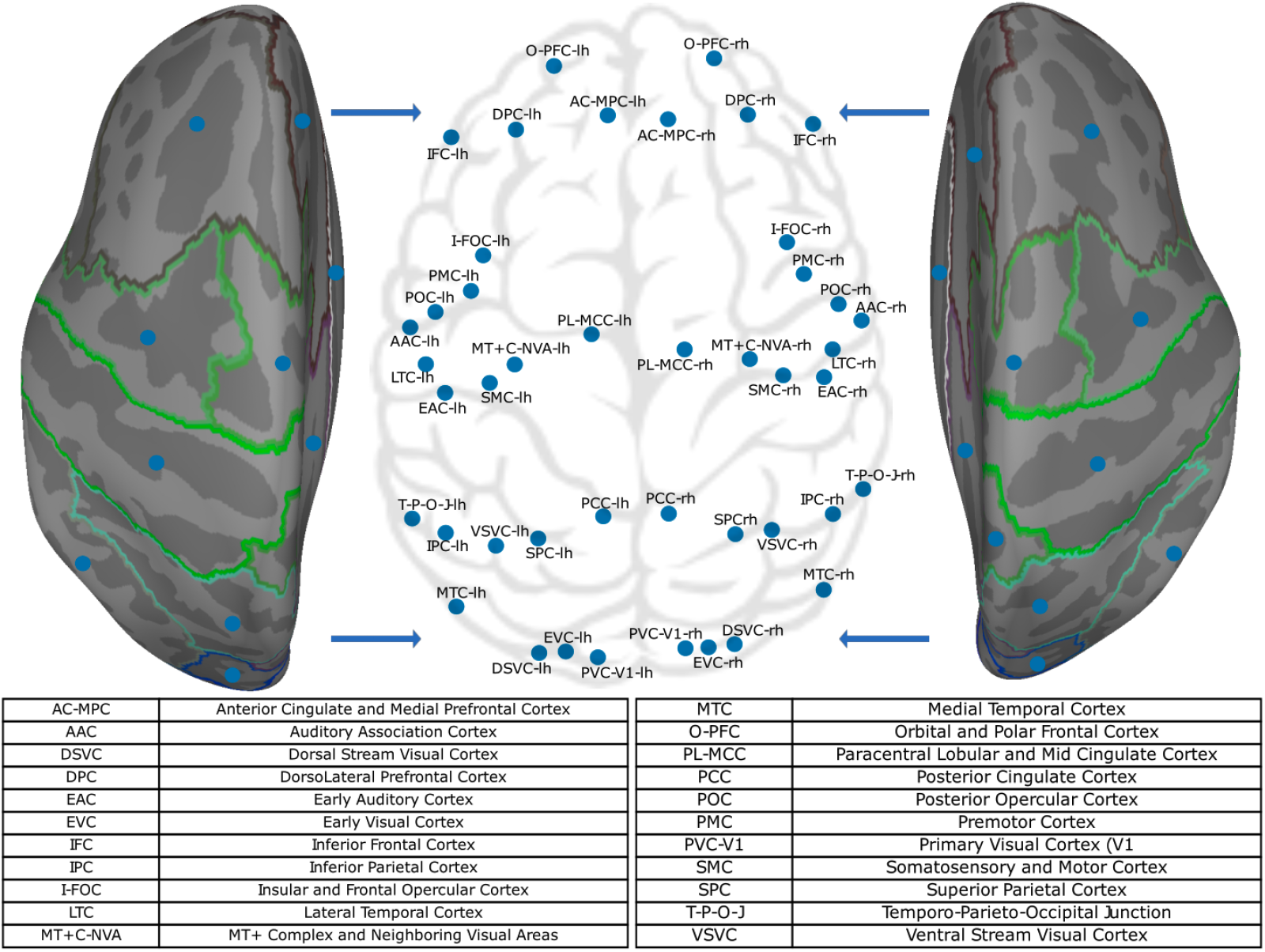
Cortex-wide projection of the centre of mass of the 44 brain regions investigated here using MEG. The brain regions were extracted using the HCP MM1 atlas.

### Extraction of the frequency peaks

For each BR of each patient we estimate the power spectral density (PSD) – using the Welch method^18^ implemented in SciPy citescipy with default parameters (average = ‘mean’ and window = ‘hann’) – averaging over 5 sec segments with 50% overlap over the whole signal. The periodic part of the spectrum was extracted using the FOOOF method proposed by Donoghue et al.^19^.

FOOOF optimally fits the PSD with a function composed of the sum of an aperiodic part and a periodic part. The aperiodic part is modelled as 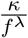 where *f* > 0 Hz is the frequency, *κ* is the offset of the PSD and λ defines how the PSD decays as a function of *f*. The periodic part is modelled as a sum of *P* weighted Gaussian distributions whose mean, standard deviation and weight respectively indicate the peak frequency, width and height of the spectral power bump of the peak. We manually specified the frequency band (1-45 Hz) on which the fitting was done. To identify oscillatory peaks in the PSD using FOOOF, we need to provide the maximum number of peaks *P*, their minimum height and band width. We searched for *P* = 4 peaks with a minimum peak height of 10^0.2^ V^2^/Hz and peak width within [1, 10] Hz (see Figure 2 C).

**Figure 2:**
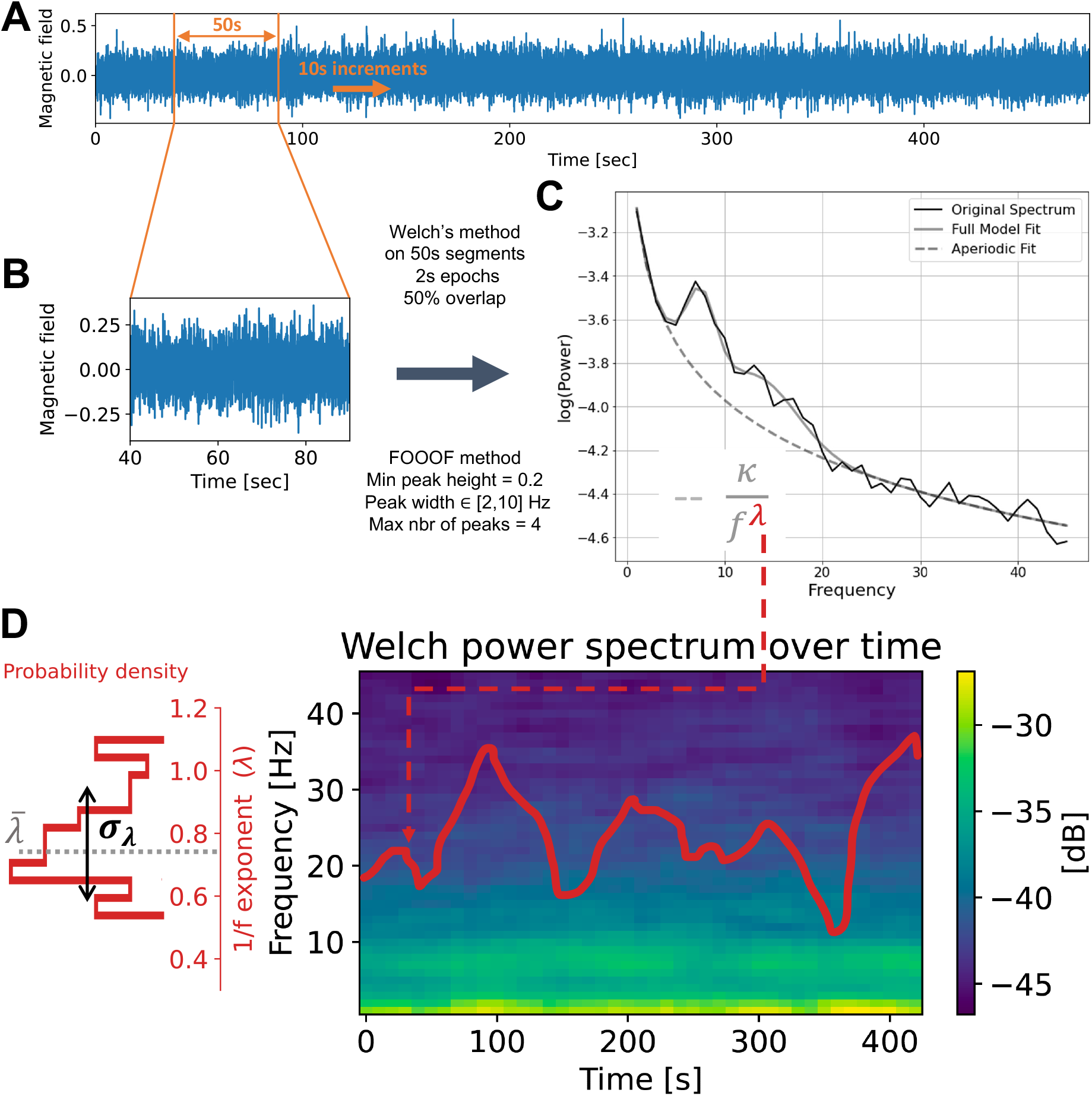
Overview of the estimation of λ dynamics done on every BR of each patient. **(A)** Time series of MEG from a source located in the pre-motor cortex left hemisphere of a PD patient. **(B)** Zoom in one epoch (duration 50 sec) of the time series of Panel A. **(C)** The black line shows the PSD of the time series of panel B. The gray line is the full fit provided by FOOOF package. The dotted line shows the aperiodic activity following a power law 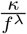. **(D)** Right: Pseudocolor image of the PSD as a function of time. The red line shows the corresponding λ for each epoch. Left: Temporal probability density of λ.

From the proportion of individuals having such peaks within each group, we defined a normalised difference between groups as

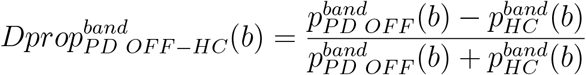

where 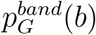 is the proportion of individual in the group *G* ∈ {*PD OFF, HC*} having at least one peak frequency in the frequency band *band* and the brain region *b* (see Figure 4 B). These proportions, 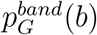, for the theta and gamma frequency bands are illustrated in Figure 4 C and D, respectively.

### Temporal dynamics of the slope of the MEG power spectrum

To characterise the aperiodic component of the MEG activity, we focused on the way the power of the MEG signals decayed as a function of the frequency. To this end, we first estimated the time-resolved spectrum (spectrogram) of each MEG source (epoch size = 50 sec, overlap 40 sec, see Figure 2 A). The PSD for each epoch was evaluated using Welch’s method averaging over 2 sec segments with 50% overlap (default parameters: average = ‘mean’ and window = ‘hann’). Next, we fitted a power law to each PSD using the FOOOF algorithm. In the aperiodic activity analysis, we used the frequency band [1, 45] Hz, a minimum peak height of 10^0.2^ V^2^/Hz, a peak width within [2, 10] Hz and a maximum of 4 peaks (see Figure 2 C for an example of such a FOOOF fitting). Finally, we obtained the temporal and spatial dynamics of *λ, κ* and *P* by applying this method to each epoch of each source (see Figure 2 D red trace). We checked the accuracy of the fit by computing its r-squared error (for all the data combined; mean: 0.94, standard deviation: 0.04).

### Calibration of λ estimate

*A priori* it is not clear how much error is expected in the estimation of λ given a specific λ and epoch size. We have fixed the epoch size to 50 sec, which is large enough to give us a good estimate of the PSD. Still we need to determine the appropriate segment size (*W*) for Welch’s algorithm. To address this issue we resorted to a numerical simulation approach. First, we created a PSD of the form 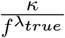, and assigned random phase (drawn from a uniform distribution 𝒰 _[−*π,π*]_) to each frequency. Next, we used inverse Fourier transform to reconstruct a signal of desired length (Figure S1 A). Finally, we extracted λ_*sim*_ for different values of *W* (Figure S1 D-F). We systematically varied λ_*true*_ and *W*. We found that the error in the estimate of λ_*sim*_ depends on both the λ_*true*_ and *W* (Figure S1 E,F). Based on these numerical experiments we chose *W* = 2 sec. which gave us the best compromise between the errors in mean and standard deviation of the λ in a range that we have observed in our data.

### Variability of λ

For each brain region, we assume that λ depends only on two parameters, time and group (PD patient ON or OFF medication and HC). So, for a brain region *b* and a subject *i*, 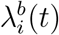 is the typical dynamics of the 1/f-exponent. To estimate the variability of 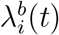 as a function of time in a given individual and BR, we calculated the coefficient of variation (CV) as:

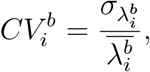

where 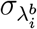 and 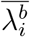 are respectively the standard deviation and the mean of λ^*b*^(*t*).

### Relationship between λ and subject age and UPDRS-III scores

λ could be related to the subject age and disease severity (UPDRS-III score) in both linear or non-linear manner. Therefore, we estimated both linear and non-linear measures of dependence between λ and *x* ∈ {*age, UPDRS*} using the following three descriptors:

- *Distance correlation* measures the non-linear dependence between two variables:

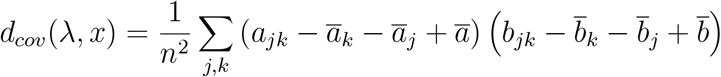

where using the notation || · || for the Euclidean norm,

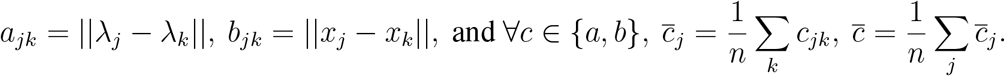

Here λ_*i*_ refers to the slope of the frequency spectrum of *i*^*th*^ subject for a given brain region. Other variables and this definition are illustrated in Figure 3. For further details please see the work by Székely et al.^20^.

**Figure 3:**
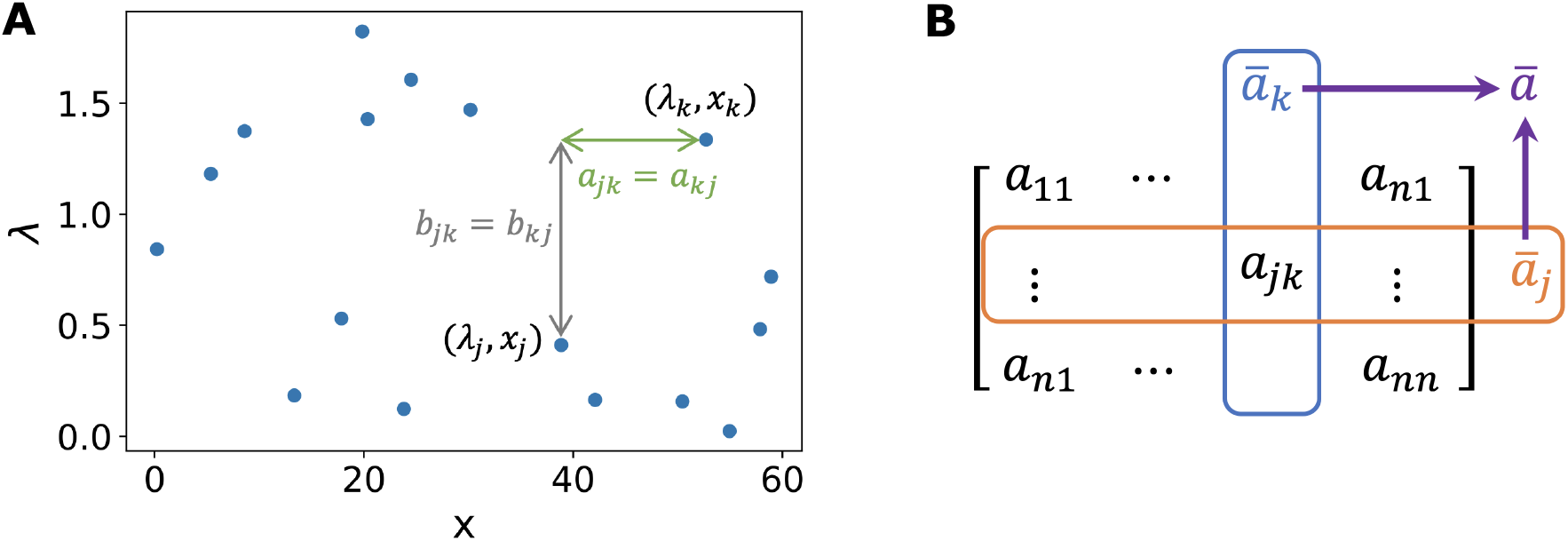
Schematic of distance correlation measurement. **(A)** Example of a sample of (*λ, x*) in a given brain region for 17 different individuals (uniformly drawn). (*a*_*ml*_)_*m,l*_ (resp. (*b*_*ml*_)_*m,l*_) is the matrix of distances within the vector λ (resp. *x*). **(B)** Computations on (*a*_*ml*_)_*m,l*_ rows and columns: *ā*_*j*_ is the mean of the *j*^*th*^ row (or column as (*ā*_*ml*_)_*m,l*_ is symmetric) and *ā* is the mean of (*ā*_*j*_)_*j*_.

- *Pearson correlation* measures the linear link between two variables:

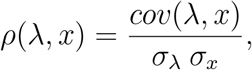

where *cov* is covariance and *σ*_*x*_ is the standard deviation of the variable *x*.
- *Spearman correlation* measures the monotonic relationship between two variables:

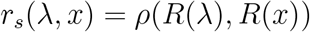

where *R*(·) is the rank function.

To disentangle the dependence of λ on the age and UPDRS-III, we computed the partial correlations. To do so, we used *partial_corr* function of the *pingouin* Python package to calculate the Pearson and Spearman partial correlations. Concerning the partial distance correlation, it is similar to the classical partial correlation. It is based on projections but now in a more complex space called the Hilbert space of U-centred matrices. We refer to the work by Székely and Rizzo^21^ for a rigorous definition. We used the corresponding function, *partial_distance_correlation*, from the Python package *dcor*^21^ for more information). We performed a permutation test (200 000 permutations on the *x* variable) to estimate the p-values.

### Statistical tests

With the exception of the correlations among UPDRS-III, age and λ, we used the Kolmogorov-Smirnov test from the *ks_2samp* function implemented in SciPy to determine the statistical significance of our results. This test captures more the deviations near the distribution centre than at its tails. Moreover, its power is greater when used in the one-tailed case. Therefore, we used the latter to test whether the cumulative distribution functions (CDFs) of one variable is greater or less in one group compared to another. In the test, the statistic is *D*^+^ = sup_*u*∈ ℝ_ [*F* (*u*) − *G*(*u*)] where *F* and *G* are the CDFs to be compared.^22^ For example, when comparing the 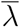 in Figure 5 B, we tested whether the CDF of 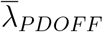 was less than 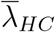 in the frontal regions and the opposite in other BRs. When comparing the two HC groups – between sessions 1 and 2, see Figure S2 B 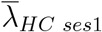 and 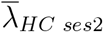 are statistically closed (distributions across the group) so that we gathered both groups as one group (the HC group). However, the CVs difference between the two sessions was of the same order than the difference between the CVs of HC and PD-OFF or PD-ON medication. Thus, we only compared CVs session-wise: PD-OFF and HC ses1, PD-ON and HC-ses2, see Figure S2 C,D. Concerning the statistical significance of correlation measures, we performed a permutation test (200 000 permutations on the *x* variable) to estimate the p-values of the partial distance correlation. For Pearson and Spearman partial correlations, two-sided p-values were computed from the one-sample *t*-test. The degrees of freedom of this test is *N* − 1 where *N* is the data size (*N* = 17 in PD groups and *N* = 20 in healthy groups).

**Figure 4:**
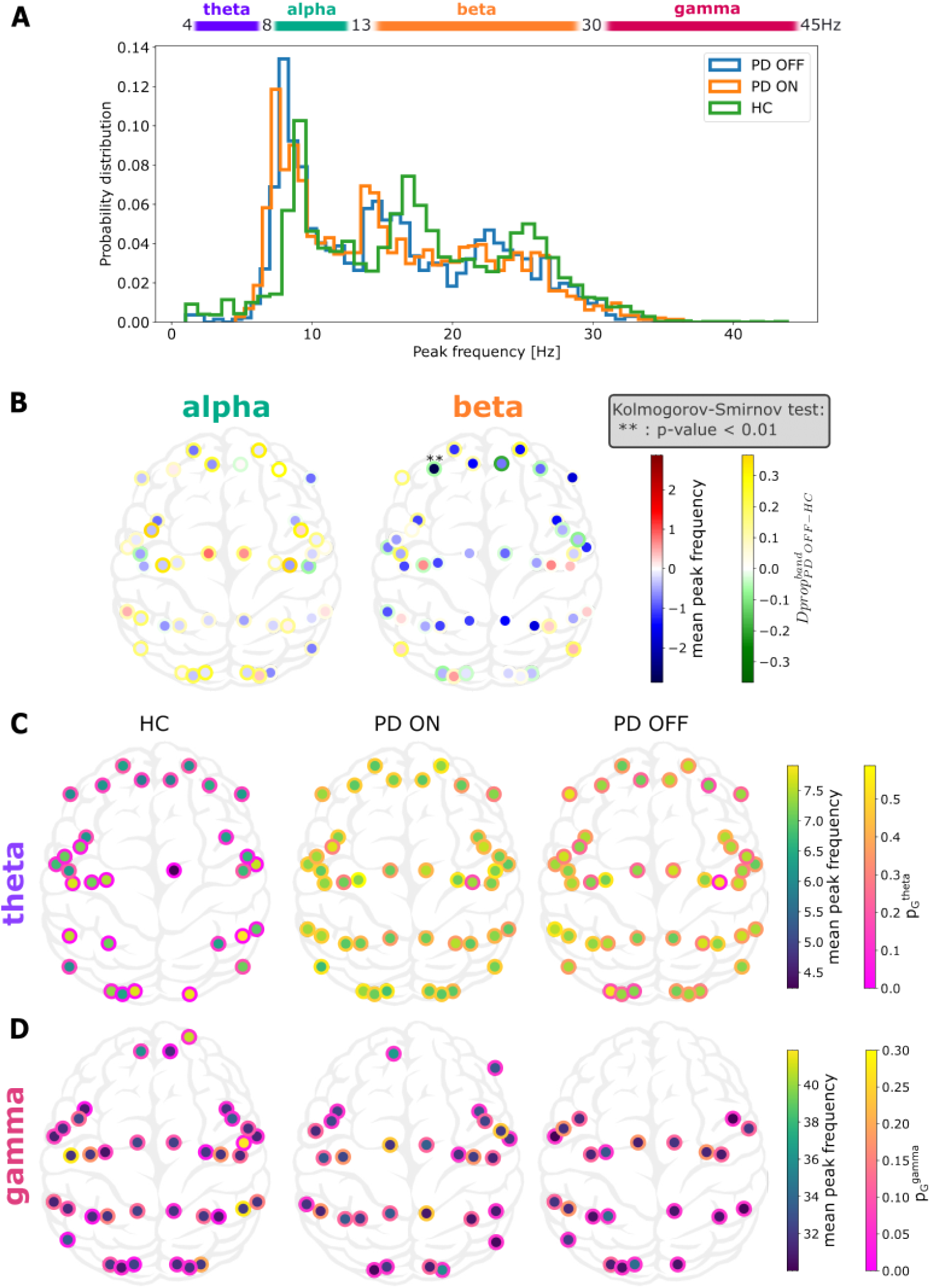
Spectral slowing, a peak frequency perspective. **(A)** Distribution of peak frequencies in all BRs of all individuals from the different groups. **(B)** Distribution of frequency peaks in alpha and beta bands. The colour inside the dots is the difference (PD-PFF - HC) of the mean peak frequency of each brain region in alpha (left) and beta (right). The edge colour of the dots is the normalised difference (PD-PFF - HC) of proportion of individuals having at least one peak frequency in the given band. **(C, D)** Distribution of frequency peaks in theta **(C)** and gamma **(CD** bands. Dot colour indicates mean peak frequency within the theta **(C)** and gamma **(D)** band. Edge colour of the dots indicate the proportion of individuals having such peaks in the given group.

**Figure 5:**
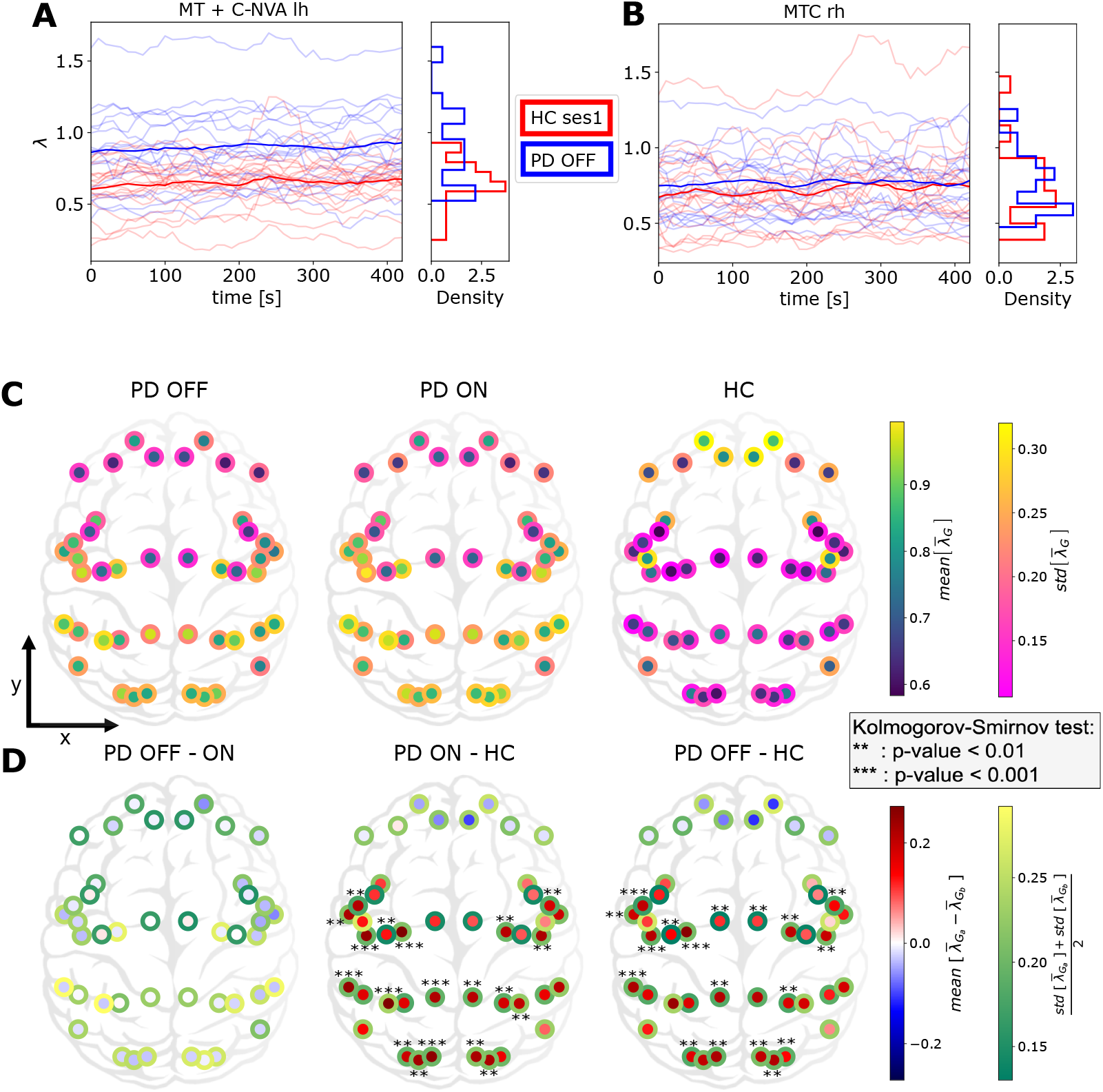
Cortex-wide distribution of the mean over time of λ within each group. **(A, B)** Temporal evolution of λ over time in two specific regions (MT+C-NVA-lh and MTC-rh) for PD patients OFF medication (pale blue) and HC (pale red). Each line corresponds to one subject. Blue and red lines indicate group averages. **(C)** Temporal average of λ_*G*_ for each group *G* ∈ {HC, PD-ON, PD-PFF}. The inside colour refers to the mean over the group. The border colour refers to the fluctuation of 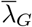 within the group *G*. **(D)** Temporal average of λ between two groups *G*_*a*_, *G*_*b*_ ∈ {HC, PD-ON, PD-PFF}. The inside colour refers to the difference between the means over the groups. The border colour refers to the averaged fluctuation of 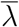 combining the two groups *G*_*a*_ and *G*_*b*_.

Finally, we did not correct for multiple comparisons. We only used comparison-wise error rate for each specific brain area, because this study is an exploratory work on the possible changes of the 1/f-exponent dynamics in PD. The alpha-level that was used to determine significance was a p-value less than 0.01. However, as we performed statistical tests on 44 BRs, a p-value of less than 0.05*/*44 ∼ 0.001 has a significance level of 0.05 after Bonferroni correction, which is the most conservative correction for multiple comparison.

## Data availability

The MEG data cannot be made publicly available because of the ethical permits. The data analysis scripts are available at: https://github.com/paschels/PD_one_over_f.

## Results

To characterise how cortical activity is changed in human PD patients, we analysed the MEG signals acquired during resting state. To this end, we quantified both the oscillatory peaks and the slope of the spectrum for MEG sources corresponding to 44 different brain regions and then we estimated the dependence of the latter on age and UPDRS-III.

### Frequency slowing in PD

Analyses of MEG and EEG from PD patients suggest that frequency associated with peak power in theta and alpha bands is reduced in human PD patients when compared to healthy controls.^3^ To test whether this is also the case in our data, we searched for Gaussian peaks in the spectrum of MEG signals without explicitly defining the frequency bands (see Materials and methods).

By estimating the distribution of all the frequency peaks observed across all the brain regions, we found that indeed in PD there is a general slowing of spectral peaks (PD-PFF vs HC, p-value < 10−20 with Kolmogorov-Smirnov test, see Figure 4 A). Notably, Levodopa medication did not improve this slowing, in fact, if at all it seemed to worsen the frequency slowing (PD-PFF vs PD-ON, p-value < 0.017 with Kolmogorov-Smirnov test).

Next, we sorted the frequency peaks into classical frequency bands (theta = 4-8 Hz, alpha = 8-13 Hz, beta = 13-30 Hz and gamma = 30-45 Hz). Thus, we obtained the proportion of peak frequency 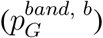 for each *band*, MEG source *b* and group *G*. Given the spectral slowing there were not enough peaks in the theta and gamma bands to compare the HC and PD groups (Figure 4 A). Therefore, first we restricted our analysis to alpha and beta bands. Even though there was a reduction in alpha band peak power, the difference did not reach a statistical significance level for none of the brain regions (Figure 4 B, left). In the beta band also we found a wide spread decrease in the mean peak frequency among most of the patients. However, the difference was statistically significant in only one brain region (Figure 4 B, right). This suggests that spectral slowing does not affect the alpha and beta bands locally. However, when we pool data across all the brain regions, spectral slowing can be observed in all bands. The main local effect (i.e. brain region specific) of spectral slowing is the reduction in the power of gamma band oscillations and increase in the power of theta bands oscillations. In particular, gamma band peaks were almost completely missing from the frontal regions in PD patients (Figure 4 D). By contrast, HC seem to be lacking theta band peaks in the central-temporal and posterior brain regions (Figure 4 C).

### Neocortex-wide change in the 1/f-exponent distribution in PD

Next, we focused on the aperiodic component of the neural activity and analysed how the spectral power decreased as a function of frequency. Using the FOOOF algorithm we fitted a power law function to the PSD (see Materials and methods) and estimated the exponent λ for each MEG source and subject. Across all the data, λ spanned a relatively wide range 0.1-2.0. However, temporal variation of λ for individual brain regions in both healthy controls and PD patients was smaller than across subject variance (Figure 5 A,B). We found that in healthy controls, λ was smaller in sensory regions than in the cognitive regions (Figure 5 C, left). However, this apparent gradient of λ from frontal to posterior regions is not statistically significant.

By contrast, in PD patients (both OFF and ON medication) λ was larger in sensory regions than in the cognitive regions (Figure 5 C, middle, right). Moreover, λ showed a clear frontal to posterior gradient (Spearman correlation, p-value *<* 10^−6^ and 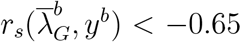 where *y*^*b*^ is the y-coordinate of the BR *b* and *G* ∈ {*PD OFF, PD ON* }).

From Figure 5 C, it is clear that there are differences in the spatial distribution of λ in PD patients and HC. To quantify this difference, we performed a brain region wise comparison of λ in HC and PD patients OFF medication (Figure 5 D right). We found that PD patients (OFF medication) have larger λ in auditory, visual, somato-senssory and motor regions than in HC (for significant figures please see Figure 5 D right). The same landscape of differences in λ was observed when we compared PD patients ON medication with HC (Figure 5 D middle). Interestingly, a comparison of PD patients in ON and OFF medication states did not reveal any significant differences across the whole neocortex (Figure 5 D left). We also note that still in most brain regions (except frontal areas), the group variance among HC was much lower than among PD patients (edge colours of the dots in Figure 5 C). Finally, the most significant changes were observed in the left hemisphere, which is expected as in our cohort most patient’s disease started in their right hand.

λ is not constant across time (Figure 5 A,B). To quantify these fluctuations over time, we measured the coefficient of variation (CV, see Materials and methods) of λ for each BR in each subject. Small value of CV of λ indicates temporal stability of λ. In general λ was stable over time (small CV). However, λ was more variable in frontal regions in both PD patients and HC (Figure S2 C). A comparison of CV of PD patients and HC revealed that for most brain regions λ was less variable in PD patients than in HC, particularly in the sensory regions where the CV difference was statistically significant (Figure S2 D,right).

Finally, even if there is no significant difference in this second order temporal statistics between PD patients ON and OFF medication, Levodopa seems to decrease temporal fluctuations in some BRs, especially frontal ones (Figure S2 D,left).

### The 1/f-exponent is positively correlated with age but not with UPDRS-III in PD

In the above we have shown how λ was altered in PD patients across the whole neocortex. The question arises: does this difference in λ also correlate with the PD severity? λ could be affected by subject age and UPDRS-III score. Therefore, we estimated partial correlations between λ and UPDRS-III given age, and age given UPDRS-III for each brain region separately.

We found that a negative partial correlation between 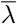 and UPDRS-III scores but these correlations do not reach a statistical significance level (Figure 6 A, see also Figure S4 for Spearman and Pearson correlations). On the other hand, in PD patients (both ON and OFF medication), 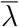 and age were positively correlated, especially in the left hemisphere T-P-O-J, SPC, PCC, MTC, MT + C-NVA, I-FOC, EAC, DSVC and LTC, VSVC in the right hemisphere (Figure 6 B, see also Table 1 for BR names, Table S1 for exact p-values and Figure S3 for Spearman and Pearson correlations). By contrast, we did not find any significant correlation between 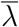 and age in healthy controls. Despite a decline in the dopaminergic neuron function due to normal ageing,^23^ these “normal” changes do not seem sufficient to affect the aperiodic component of cortical population activity, unlike in PD.

**Figure 6:**
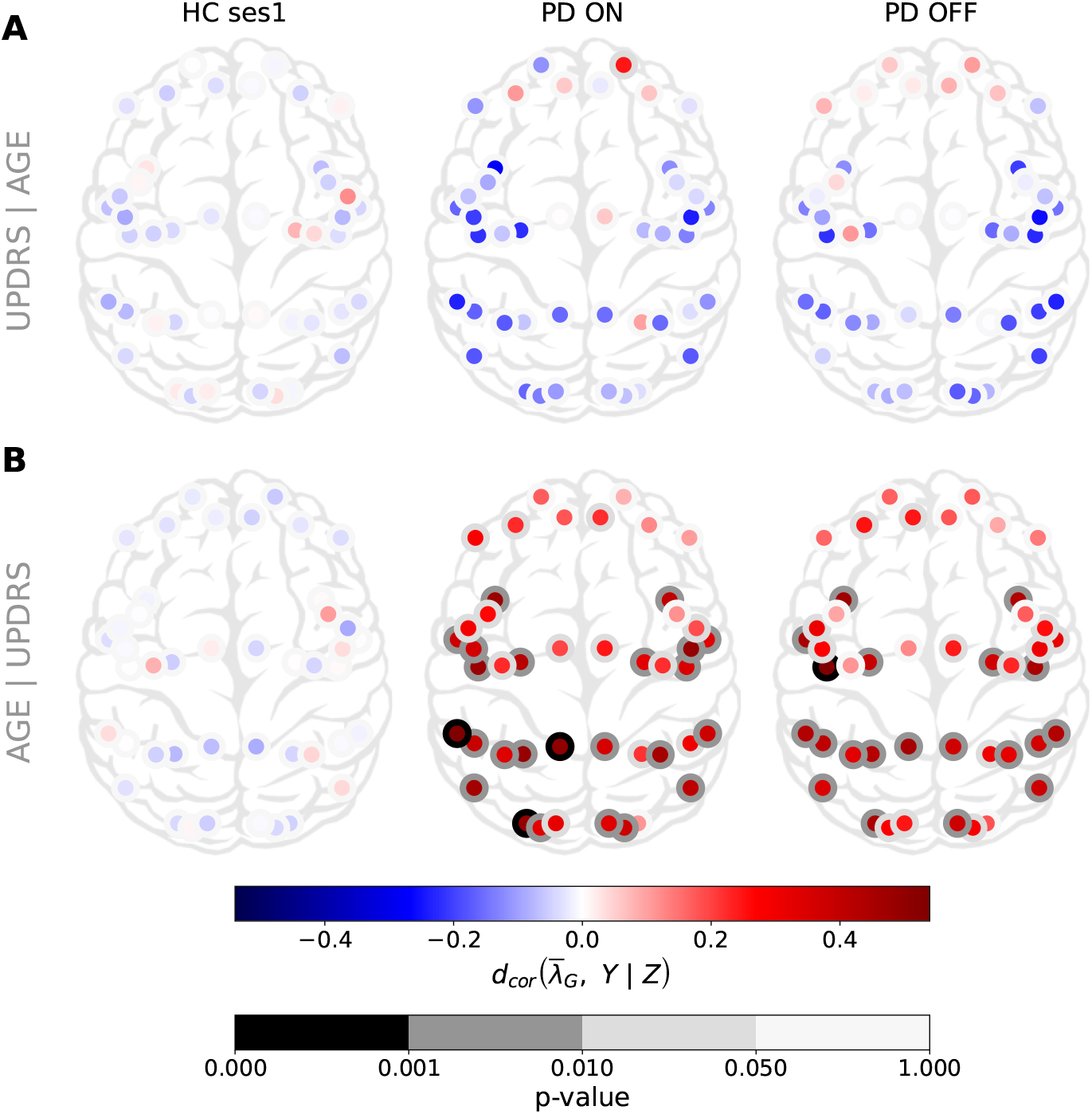
Cortex-wide distribution of partial distance correlation between the mean over time of 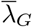 and *Y* knowing *Z* for each group. *G* ∈ {*HC ses*1, *PD ON, PD OFF* }. **(A)** (*Y, Z*) = (UPDRS,age). **(B)** (*Y, Z*) = (age,UPDRS).

It is somewhat counter intuitive that despite significant PD-related changes in 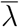, it was not correlated with UPDRS-III, even in the motor regions of the cortex. This result may be explained by a lack of sensitivity of the data or by the fact that 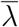 is not by itself related to motor symptoms. The fact that the increase in 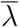 was positively correlated with age suggests that persistent dopamine depletion may be more detrimental for brain networks that are already undergoing age related deterioration like the sensory areas. For instance, vision is impaired in PD, especially in patients with cognitive decline.^24,25^ Alternatively, brain networks in the younger patients may be more malleable to compensate for the loss of dopamine than in older patients.

## Discussion

Given chronic change in the neuromodulator dopamine we expect that excitation and inhibition (EI) balance may be altered in the neocortical regions. To test we analysed the aperiodic activity of MEG data which has been inversely related to EI balance in several previous studies.^26–35^ We found that the 1/f-exponent (λ) was higher in PD patients than in HC in most brain regions but not in frontal regions. The most significant changes occurred within sensory and motor regions. Moreover, we found that λ showed a spatial gradient from posterior to anterior brain regions in PD patients. Although not significant, in HC the gradient of λ was reversed compared to that in the PD patients. When examining the fluctuations of λ over time, we found that λ was more variable in the frontal regions. Moreover, λ fluctuated more in HC controls than in PD patients. This was not expected as λ was globally larger in PD. Surprisingly, l-dopa medication had a little effect on the topography of λ. Finally, we computed different correlation measures between λ and both the age and UPDRS-III score. We did not find any correlations in the HC group neither with age or UPDRS-III. Although not significant, global negative correlations with UPDRS-III were observed in PD patients. On the other hand, λ was positively correlated with age in PD patients in both ON and OFF medication states.

Previously, Vinding et al.^36^ analysed the same data for 1/f slope (λ) but that analysis was restricted to the sensorimotor association regions. Consistent with our results, they also found that λ was positively correlated with age in PD patients but not in HC and the correlation between λ and UPDRS-III was not significant. We build on that study and now show the topography of λ over the whole neocortex in both healthy controls and PD patients. Our analysis shows that λ changes in early sensory regions are also correlated with age but only in PD patients.

Recently, Wang et al.^13^ analysed the 1/f slope of EEG activity. Our results differ from their results in several ways. First, concerning the topography of λ’s topography, we found that λ decreased (increased) from sensory to cognitive regions in PD patients (healthy controls). By contrast, Wang et al.^13^ have reported similar topography in both healthy and PD patients. In their study, λ was high in the central regions with a decrease in all directions from there on. This difference might be due to the fact that we used a brain atlas to project the MEG sensors to sources. Moreover, we have a much denser spatial sampling of the space using 306 MEG sensors as opposed to the 32 EEG sensors used by Wang et al.^13^. However, this is a conspicuous difference that should be investigated in a study where both MEG and EEG are acquired from the same patients. In contrast to Wang et al.^13^, we did not find significant differences between PD-ON and PD-OFF states. This could be due to the difference in recording protocols in the two studies: in our case, dopamine effects were acute in the sense that patients were OFF medication for at least 12 hours but then took the medication only one hour before the ON medication state was recorded. Unlike Wang et al.^13^, we have analysed partial correlation between UPDRS-III/age and each MEG source. This separation of brain regions and partial correlation revealed that λ was correlated to age but only in PD patients.

### Interpretation of 1/f slope

There are at least three possible interpretations of the changes in λ. Population signals such as MEG are generated by dipoles created by transmembrane currents.^37^ The transmembrane currents are generated due to synaptic inputs impinging on the neurons. Therefore, the frequency spectrum of the MEG (and also EEG, LFP) reflects the time constants of the synaptic inputs. Typically, inhibitory (GABAergic) synaptic transients have a longer time constant than excitatory (glutamatergic) current. Therefore, increased λ may be an indicator of either increased GABAergic or reduced glutamatergic currents. That is, λ is a proxy of relative excitation-inhibition (EI) balance.^10^

However, even glutamatergic synaptic inputs due to NMDA receptors can also be very slow. Therefore, it is also possible that increased λ may indicate a relative increase in the NMDA type synaptic currents. In the context of our results, we found that λ in PD is increased in the sensory areas. These areas usually have relatively small fractions of NMDA receptors.^38^

Finally, it is also possible that the change in λ is simply a reflection of changes in the dynamics of local and external inputs to the network. That is, if the network is driven by slowly fluctuating inputs it will also reflect in slower fluctuations and therefore larger λ. If this is the case then we would also expect a change in the time scales of spiking activity.

### Hypotheses about network level changes in PD

Each of these possibilities suggest a different but testable hypothesis. If the slope of the MEG reflects change in the relative fraction of excitatory and inhibitory currents, then it is tempting to hypothesise that in PD there is an increase in inhibition in the sensory region. Increase in relative fraction of NMDA will also account for an altered excitation-inhibition balance on longer time scales. Techniques such as MRS could be used for a non-invasive estimate of the relative fraction of AMPA, NMDA and GABA in order to test this hypothesis. Ideally, these hypotheses should be tested in animal models with *in vivo* measurements of excitatory and inhibitory currents.

If changes in the λ reflect a change in the time scales of fluctuations in the input then, we will need to explain how in a presynaptic network the spectrum of neuronal activity could change. For that we again revert to the hypothesis of a change in excitation-inhibition balance. However, before following this line of reasoning, we need to estimate whether there are significant changes in the spiking activity of a given brain region where λ has changed. This suggests that spiking activity from early sensory regions should also be recorded in animal models of PD.

As we discussed earlier, it is tempting to relate the slope of the frequency spectrum to EI balance. Origin of pathological activity in the basal ganglia during PD, especially the beta band oscillations is closely related to changes in the EI balance in STN and GPe regions of the basal ganglia.^39,40^ Local field potential recorded from the STN or GPe during DBS surgery can be used to estimate relative EI balance at different locations in the STN in terms of λ s. Such an estimate of relative EI balance could guide stimulation electrode placement.

### Age and UPDRS-III dependence

A lack of a clear correlation between *λ s* and UPDRS-III suggests that it may not be useful as a clinical biomarker. However, correlation between *λ s* and age in the PD group suggests that λ is indeed altered in PD. In particular, we observe a positive correlation in PD in most brain regions except the frontal regions. This result may suggest that dopamine depletion has an important impact on the sensory and motor ‘ageing’.

A positive correlation between λ and age in PD patients is a curious observation. Previous work from several groups have shown a negative correlation between λ and age in the neocortex.^41–43^ However, in over 60 years old HC, λ from somatosensory regions may positively correlate.^44^ In contrast to these findings, in our data we did not observe any significant correlation between λ and age in HC. As age is highly (positively) correlated to disease duration, PD thus seems to affect all (except frontal parts) necortex’s *λ s* by increasing them over age/disease duration.

### Effect of dopamine on λ

Levodopa effects are fast as shown by the UPDRS-III score improvements only one hour after the drug administration. We found that in our data Levodopa influenced the spectral slowing. However, the effect of Levodopa on λ was not observed in the neocortex. To the best of our knowledge, there is no consensus on the cortical effects of Levodopa.^36^ In our study, a possible reason could be the short time (1 hour) between taking the medication and recording of MEG. A recent study by Wang et al.^13^ reported changes in the λ estimated from EEG. In that study, PD patient ON medication took their medication dose as usual, in the morning before the measurement. However, Wang et al.^13^ also reported that Levodopa did not improve the aperiodic part (i.e. λ increased) of the neocortex activity spectrum. In addition, the total washout of Levodopa may take days.^45^ The latter could explain the similar results we obtained for ON and OFF medication.

## Limitations

Here we used a commonly used range of frequencies (1 to 45 Hz) for our analysis. This range is relatively small. Indeed the broader the frequency range, the better the fit should be. However, for MEG it is not as easy to take the largest band possible because muscle artifacts become more prominent in high frequencies. Therefore, local field potential or ECoG may be more suited for such an analysis over broad frequency ranges.

Next, we have used a specific brain atlas and it is not clear how the topography of λ may change when using a different brain atlas. In this regard, adding information from the sensor space may help matching the different atlases’ results.

Finally, to better understand the functional implications of λ changes, a more detailed correlation analysis is needed that takes into account the UPDRS-III sub-scale^46^ as well as cognitive scores.

### Implications

Our findings also raise the question why frontal regions are more protected than sensory and motor regions even though dopaminergic projections in the neocortex are primarily restricted to the frontal regions.^47^ The lack of prefrontal changes between PD and HC may be due to dopamine pathways. Indeed, the main brain area producing dopamine and projecting to the cortex (in particular prefrontal cortex) is the Ventral Tegmental Area, which is altered after the substantia nigra compacta in PD.^48^ Hence, changes in the frontal regions may take longer to manifest.

In addition, because there is such a big change in the λ value in sensory regions one would expect deficits in the sensory representations. It is well established that PD patients have olfactory,^49^ proprioceptive^50^ and cross-modal sensory fusion deficits.^51^However, our work suggest changes in other sensory modalities such as vision and audition.

It is common to measure functional connectivity between brain regions in a frequency dependent manner. Usually these estimates are based on filtered time series. Here we found that λ varies over time. Therefore, we can ask whether λ variations across brain regions are correlated or not in PD and HCs. Recent work suggests that functional connectivity based on the component of aperiodic activity may be more robust.^52^

Overall, we show that the aperiodic activity, which usually has been considered as noise, gives new insights in PD and deserves more attention when analysing any neural field potentials like ECoG, LFP, EEG and MEG. Finally, previous studies showed frequency slowing in relation to cognitive decline rather than the motor symptoms.^5^ It could then be that the change in λ is a more general expression of neurodegeneration than only the dopamine affected systems.

## Abbreviations

PD: Parkinson’s disease
BR: brain region
MDS-UPDRS: Movement Disorder Society’s Unified Parkinson’s Disease Rating Scale
EI: excitation-inhibition
BG: basal ganglai
FC: functional connectivity
HC: healthy control
PSD: power spectral density

## Acknowledgements

We thank Archishman Biswas for his help on the Figure S1.

## Supplementary Material

**Figure S1:**
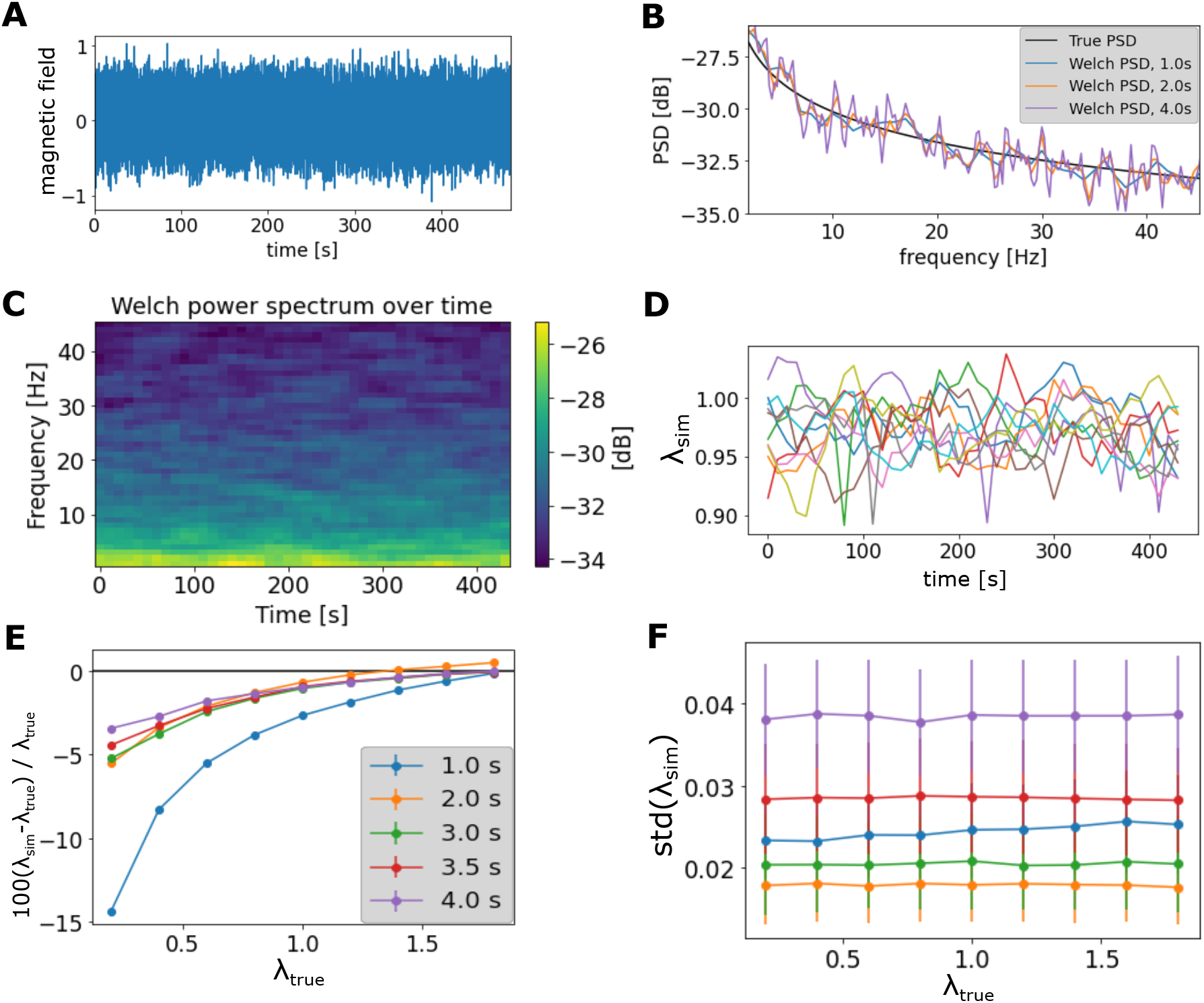
Comparing the true 1/f-exponent of a simulated signal reconstructed from a given power spectrum of the form *κ/f*^λ^. **(A)** Reconstructed signal from PSD equals to 10^*−*2.5^*/f* ^0.5^. **(B)** PSD of the reconstructed signal using the entire signal or a segment of it of sizes 1, 2 or 4 sec. **(C)** Spectrogram of the reconstructed signal shown in **A** using the method described in Materials and methods. **(D)** Temporal dynamics of λ_*sim*_ the reconstructed signals for a PSD 10^*−*2.5^*/f* ^1^ and epochs of 1 sec. **(E)** Temporal average of λ from reconstructed signals, λ_*sim*_, compared to the true λ_*true*_. **(F)** Temporal standard deviation of λ from reconstructed signals in function of the true λ.

**Figure S2:**
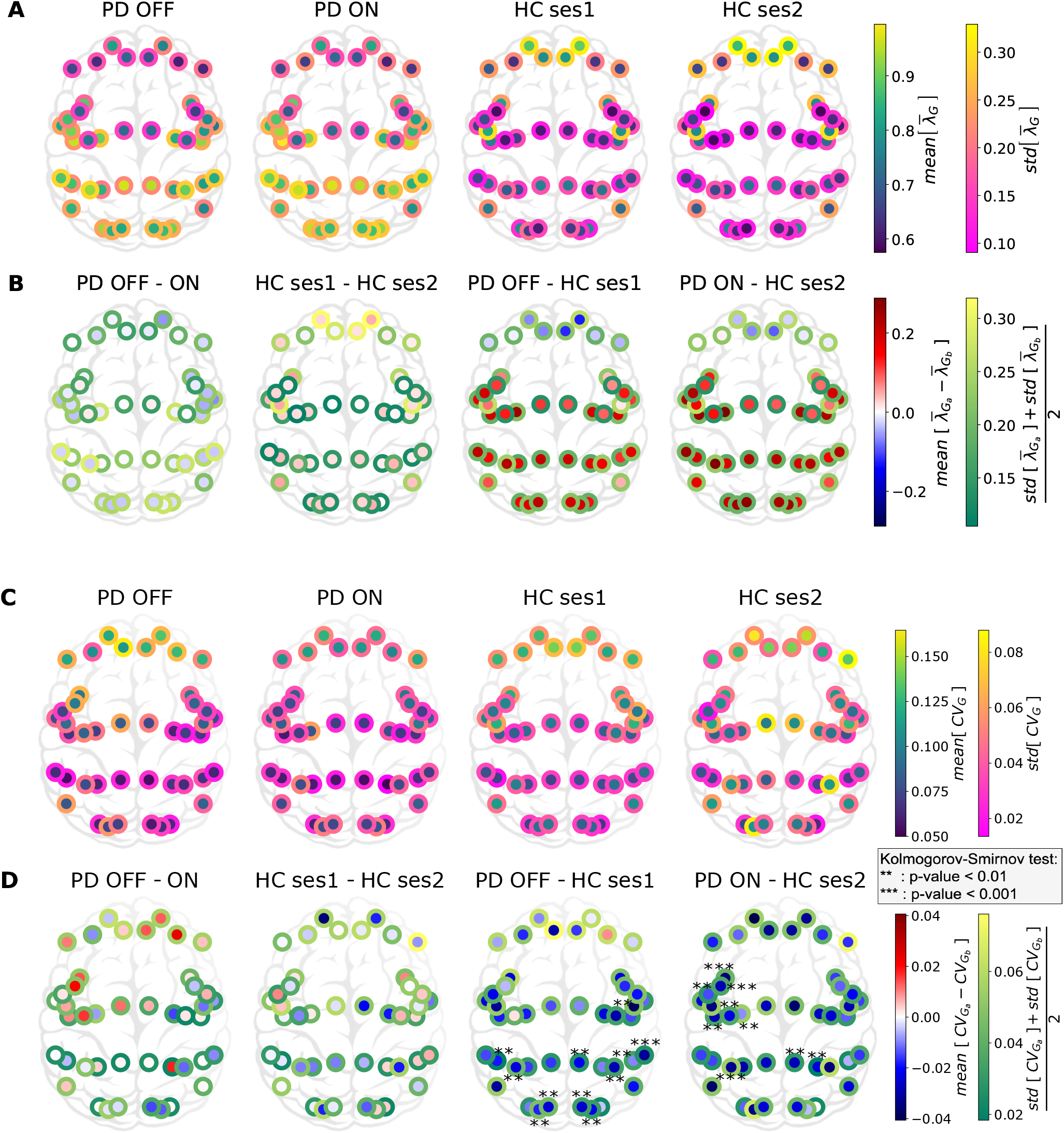
Cortex-wide distribution of the mean and the coefficient of variation over time of λ within each group. **(A)** (resp. **(C)**) Temporal mean (resp. coefficient of variation) of λ_*G*_ for each group *G* ∈ {HC ses1, HC ses2, PD-ON, PD-PFF}. The inside colour refers to the mean over the group. The border colour refers to the fluctuation of 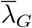 (resp. *CV*_λ*G*_) within the group *G*. **(B)** (resp. **(D)**) Temporal mean (resp. coefficient of variation) of λ between two groups *G*_*a*_, *G*_*b*_ ∈ {HC ses1, HC ses2, PD-ON, PD-PFF}. The inside colour refers to the difference between the means over the groups. The border colour refers to the averaged fluctuation of 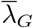 (resp. *CV*_λ*G*_) combining the two groups *G*_*a*_ and *G*_*b*_.

**Figure S3:**
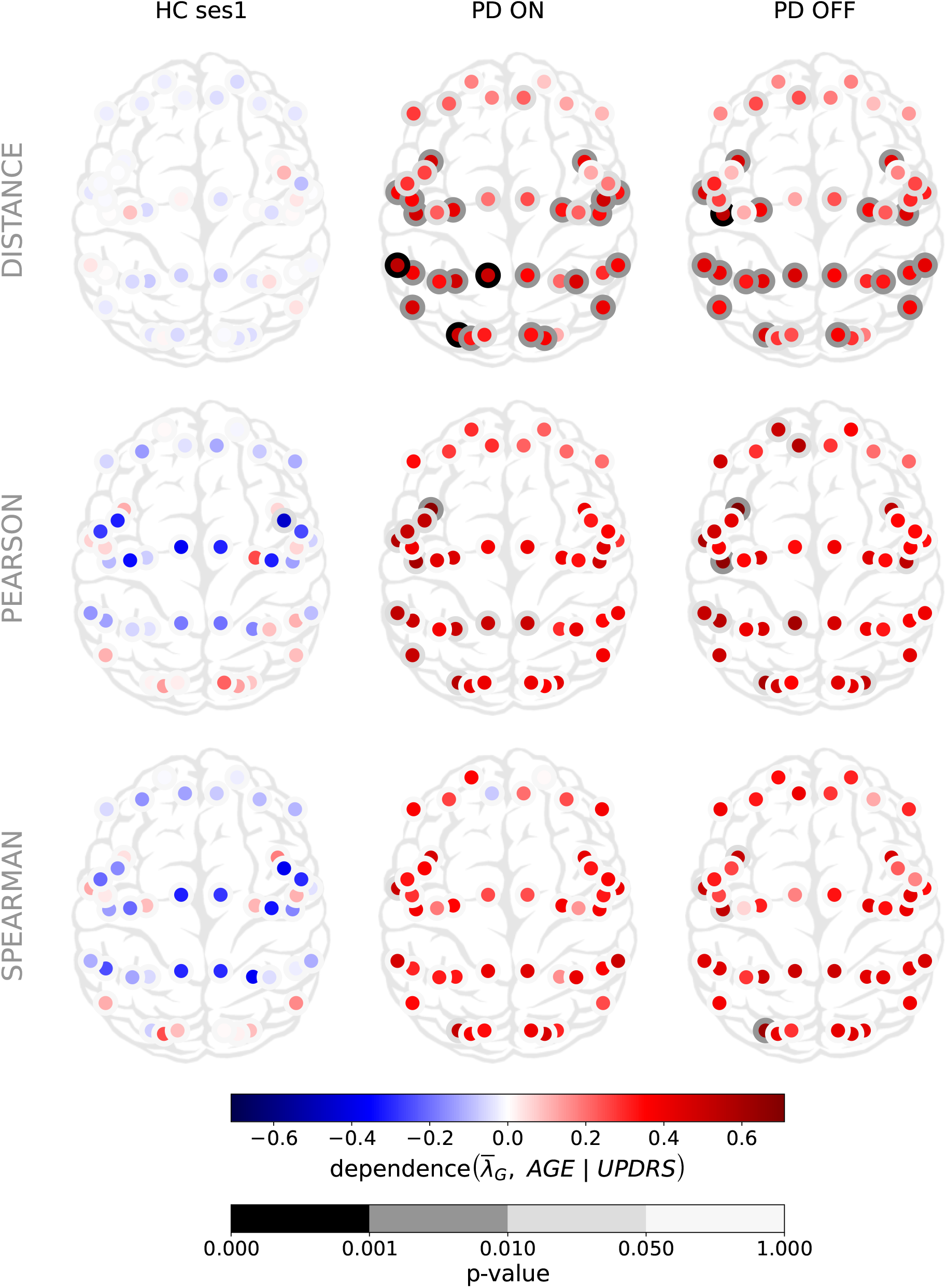
Cortex-wide distribution of different relationship measures between the mean over time of λ and ages within each group.

**Figure S4:**
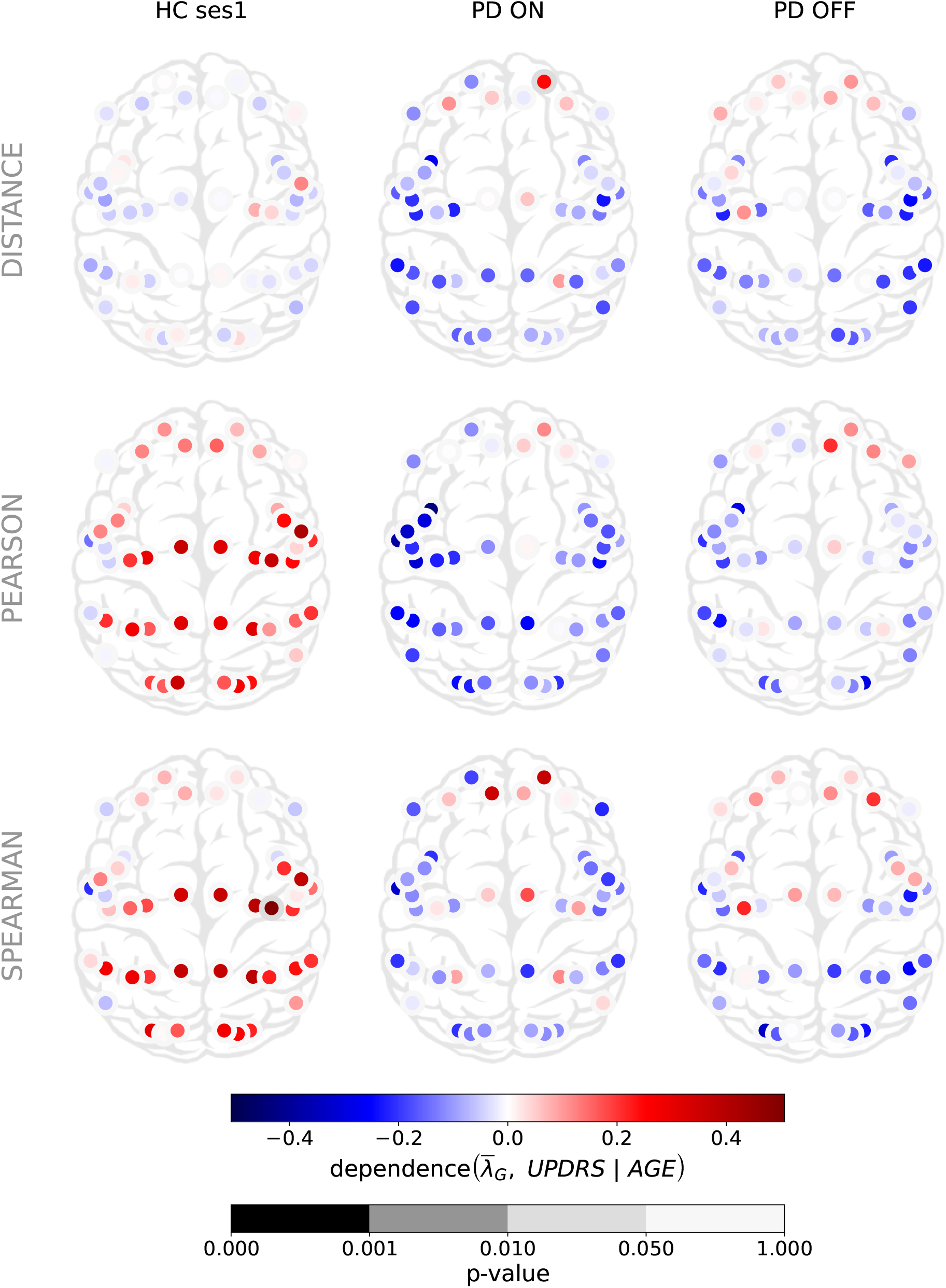
Cortex-wide distribution of different relationship measures between the mean over time of λ and UPDRS-III within each PD group and combining them (first column).

**Figure S5:**
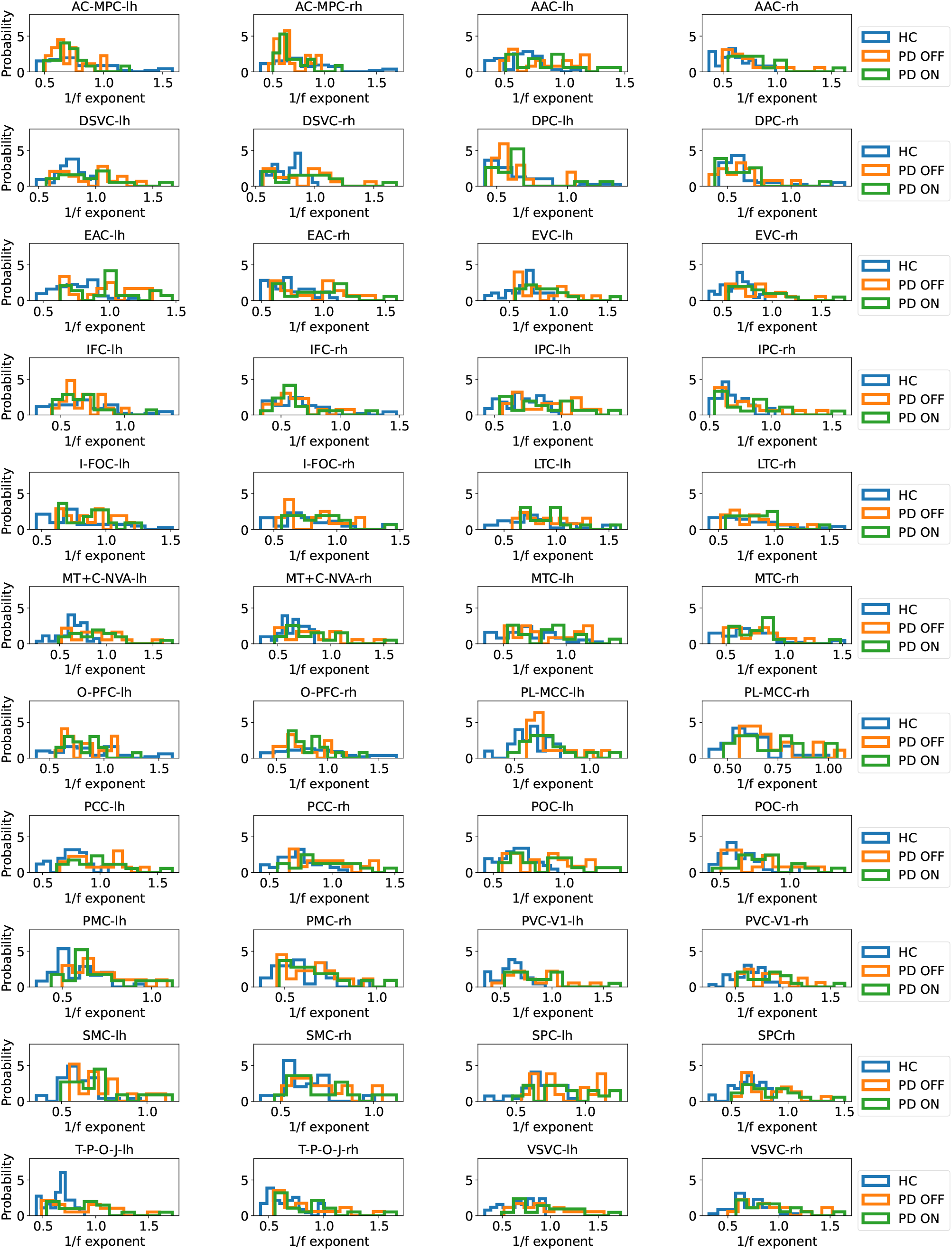
1/f-exponent distributions over the different groups in the different BRs.

**Figure S6:**
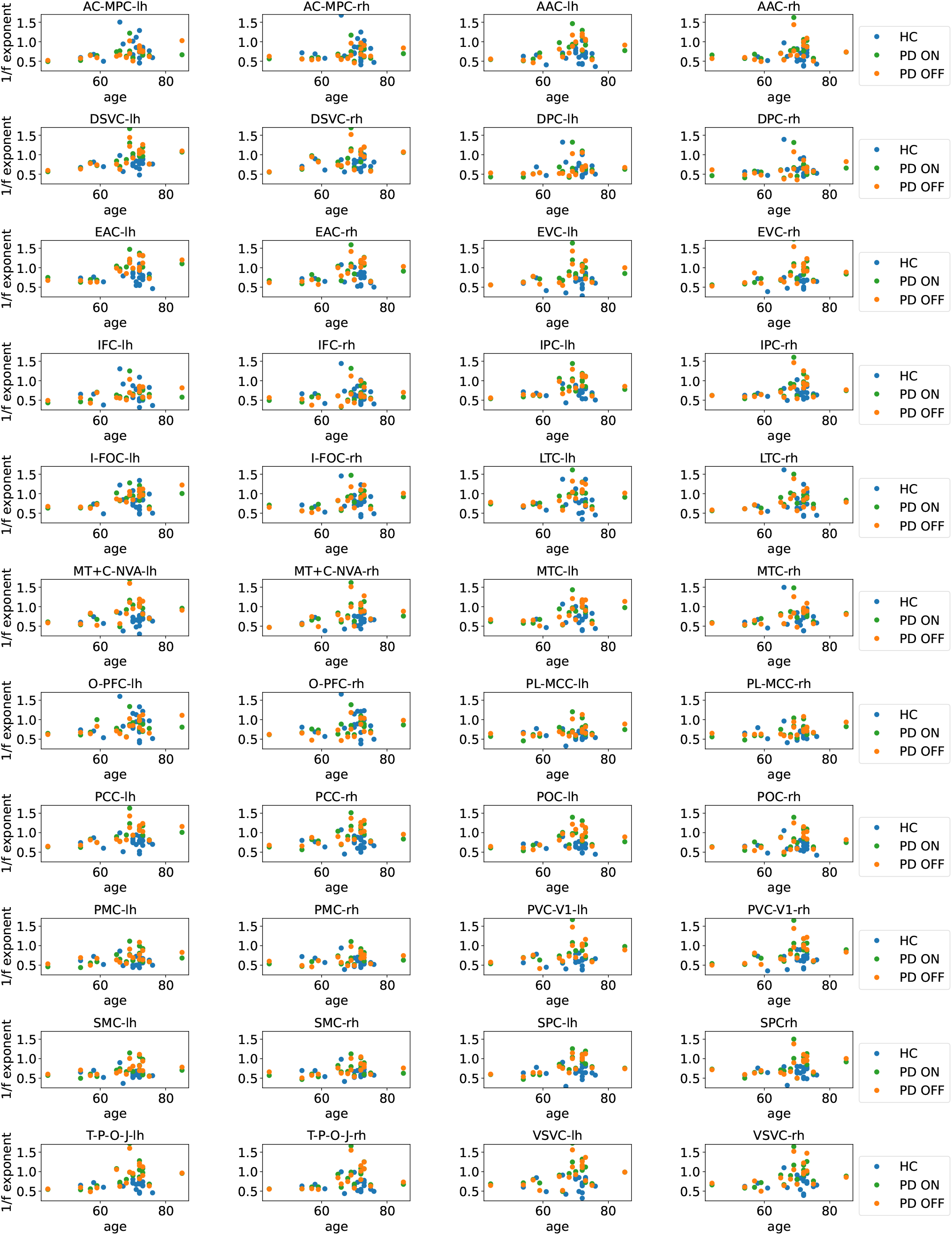
1/f-exponent as a function of age for the different groups in the different BRs.

**Figure S7:**
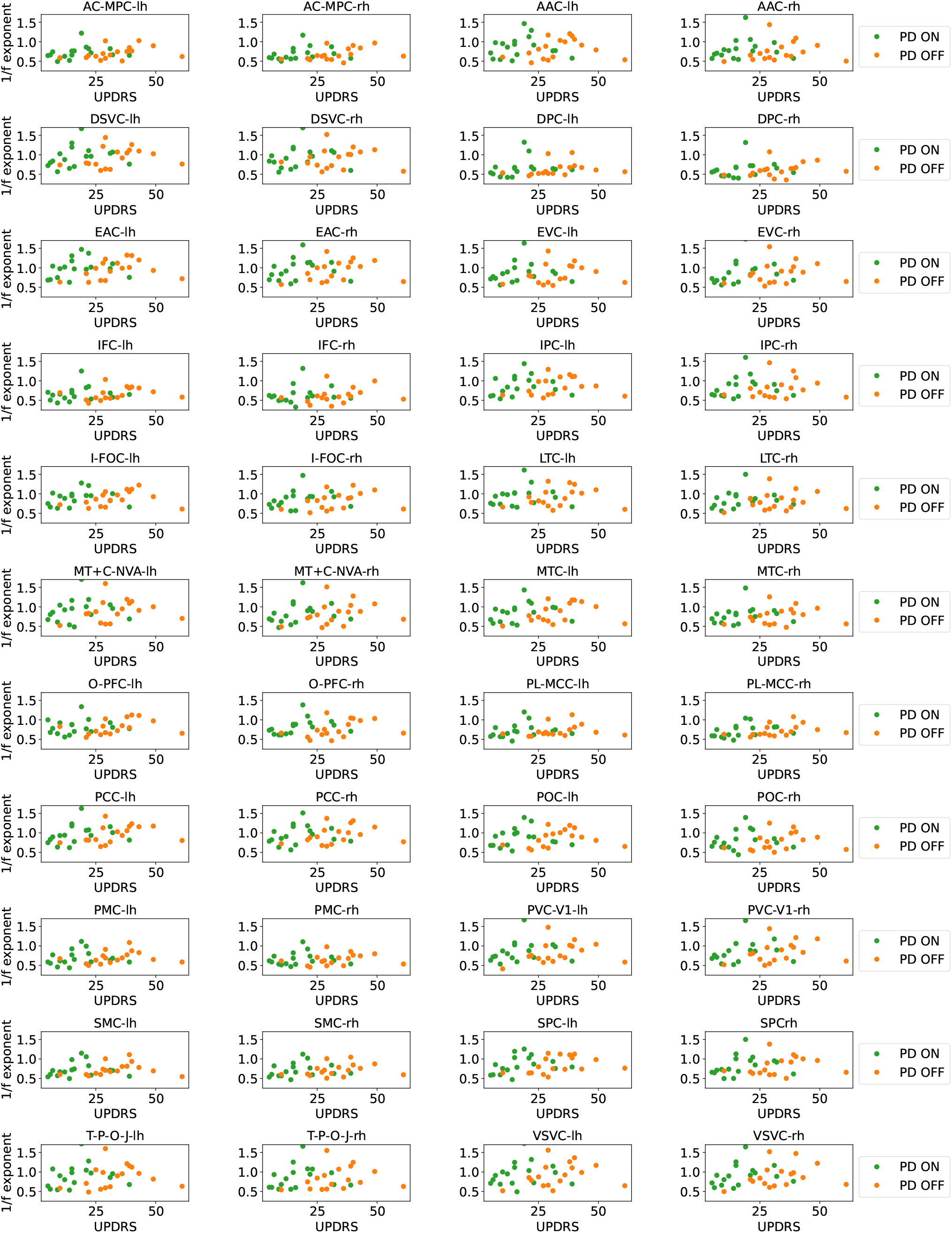
1/f-exponent as a function of UPDRS-III for PD-ON and PD-OFF in the different BRs.

**Table S1:** P-values of the distance correlation per brain region, PD patient session and hemisphere. The shadowed cells are the 20 lowest p-values.

| Patient<br>Brain region | PD ON lh | PD ON rh | PD OFF lh | PD OFF rh |
| --- | --- | --- | --- | --- |
| AC-MPC | 0,052655 | 0,0327 | 0,023435 | 0,05004 |
| AAC | 0,005475 | 0,008165 | 0,001995 | 0,01079 |
| DSVC | 0,00098 | 0,10805 | 0,001665 | 0,05319 |
| DPC | 0,029015 | 0,09365 | 0,020805 | 0,138505 |
| EAC | 0,00134 | 0,00813 | 0,00047 | 0,002055 |
| EVC | 0,009255 | 0,00509 | 0,017115 | 0,016085 |
| IFC | 0,0169 | 0,11816 | 0,065705 | 0,072345 |
| IPC | 0,004815 | 0,014195 | 0,00435 | 0,006125 |
| I-FOC | 0,001375 | 0,00364 | 0,00146 | 0,002615 |
| LTC | 0,00621 | 0,00102 | 0,0191 | 0,01329 |
| MT+C-NVA | 0,00308 | 0,0098 | 0,004235 | 0,00613 |
| MTC | 0,001795 | 0,004075 | 0,008265 | 0,00591 |
| O-PFC | 0,05662 | 0,15312 | 0,061315 | 0,053295 |
| PL-MCC | 0,0401 | 0,02616 | 0,09309 | 0,02303 |
| PCC | 0,00078 | 0,007705 | 0,00162 | 0,006045 |
| POC | 0,015555 | 0,04716 | 0,012155 | 0,02366 |
| PMC | 0,018335 | 0,100665 | 0,12863 | 0,05784 |
| PVC-V1 | 0,01139 | 0,00877 | 0,024695 | 0,00548 |
| SMC | 0,025775 | 0,033375 | 0,10384 | 0,02987 |
| SPC | 0,001025 | 0,0362 | 0,003105 | 0,014585 |
| T-P-O-J | 0,000495 | 0,006065 | 0,002675 | 0,0031 |
| VSVC | 0,00903 | 0,001855 | 0,00938 | 0,009635 |

